# Re-evaluating the eukaryotic Tree of Life with independent phylogenomic data

**DOI:** 10.64898/2026.04.08.717153

**Authors:** Romain B. Leroy, Laura Eme, Purificación López-García, David Moreira

## Abstract

Understanding the phylogenetic relationships among eukaryotic lineages is essential for tracing the evolution of key phenotypic traits and inferring the nature of the Last Eukaryotic Common Ancestor. While phylogenomic analyses have clustered eukaryotic taxa into several well-supported major ‘supergroups’, the relationships among them remain largely uncertain. Phylogenetic signal erosion over deep time and limited available taxon sampling are among the possible causes. However, most previous studies rely on variations of the same core protein dataset, hence containing the same potential systematic biases. Here, we reconstructed the eukaryotic Tree of Life using a largely independent, marker-rich dataset derived from highly conserved Benchmarking Universal Single-Copy Orthologs. Unlike previous collections, our 277-marker supermatrix minimizes ribosomal protein representation and shares less than 25% overlap with previous datasets. State-of-the-art analyses of this dataset confirm most eukaryotic supergroups previously observed, but suggest different positions for some lineages. Notably, Telonemia clusters with Haptophyta rather than SAR (Stramenopiles-Alveolata-Rhizaria), and Ancyromonadida and Malawimonadida form a monophyletic group at the base of the Opimoda. Our results highlight the importance of analyzing independent phylogenomic datasets and support the hypothesis that extant eukaryotic diversity encompasses a small number of high-rank, supergroup lineages.

## Introduction

Eukaryotes, defined by the presence of a nucleus, typically possess numerous other complex intracellular features, such as mitochondria, a dynamic cytoskeleton, and an intricate endomembrane system. While they include nearly all macroscopic life forms (animals, fungi, plants, and macroalgae), the vast majority of their diversity lies among the myriad lineages of unicellular species, generally referred to as protists, which exhibit an extraordinary range of cell architectures, lifestyles, and genetic diversity^1^. Understanding the evolutionary relationships between major eukaryotic phyla remains a central challenge in evolutionary biology^2–5^. A resolved eukaryotic Tree of Life (eToL) offers the necessary framework to retrace how eukaryotes diversified into current lineages from their ∼2-billion-years-old Last Eukaryotic Common Ancestor (LECA) and map the evolution of defining eukaryotic traits^6^.

Most eukaryotic phyla appear to have diversified over a relatively short evolutionary timespan, a process referred to as a radiation^7^. As a result, phylogenetic distances among them are very short, making their relationships difficult to infer and requiring large amounts of data to get resolved. Taking advantage of the increasing amount of genome and transcriptome sequences of diverse eukaryotes^8^, this challenge has been addressed using phylogenomic (i.e., multi-gene) analyses over the past few decades^4,9–17^. These analyses have consistently shown that most eukaryotic lineages cluster into a handful of ‘supergroups’, which constitute the current paradigm to depict the organization of all eukaryotic diversity^1,18^. The most solidly established ones include SAR (Stramenopiles, Alveolata, and Rhizaria) and Amorphea (Amoebozoa, Opisthokonta, Breviata, and Apusomonadida)^1,11,14^. Other classical supergroups have been less consistently supported across analyses. This is the case of Archaeplastida, composed of plants and algae with primary plastids but including other groups, such as cryptophytes, in some analyses^4,5,19–21^, and Excavata, a historical assemblage of diverse biflagellates based on a shared morphological trait: a ventral feeding groove hosting the posterior flagellum^4,5,19–21^. In addition to these major supergroups, several other smaller lineages (such as Ancyromonadida and Apusomonadida^4^, CRuMs^15,22^, Malawimonadida^20^, Haptista and Cryptista^21^, or the recently described predatory phylum Provora^23^), complete the known diversity of eukaryotes. Although the deep relationships among these groups remain uncertain, they appear to join the previously cited supergroups into even larger lineages (‘megagroups’), such as the Opimoda (Amorphea, CRuMs, Malawimonadida, and Ancyromonadida) and the Diaphoretickes (SAR, Archaeplastida, Haptista, Telonemia, and Cryptista)^1,24^, with a major apparent split in the eukaryotic tree occurring between Opimoda and Diphoda^4,5,25^.

Several protein marker collections have been used to infer these phylogenetic relationships. Importantly, these phylogenomic datasets mostly derive from one initial study^9^ (Supplementary Fig. 1), largely reused in subsequent analyses. While this approach has allowed the reconstruction of a relatively stable phylogeny for many eukaryotic lineages over time, it may also have perpetuated potentially biased phylogenetic signals contained in these proteins. Therefore, it becomes crucial to compare previous results with those based on independent sets of markers to rule out the possibility that the supergroup structure of the eToL contains artefacts, and to improve the resolution of the phylogenetic relationships of still-unstable lineages.

One untapped source of highly-conserved protein markers is the Benchmarking Universal Single-Copy Orthologs (BUSCO) dataset, a collection of essential, near-universal single-copy genes widely used to quantify genome and transcriptome completeness^26^. By definition, BUSCO genes are found across virtually all eukaryotes, making them attractive candidates for large-scale phylogenomic analyses^27^. In fact, recent studies have begun to explore their utility to resolve evolutionary relationships within specific clades of plants or animals^28,29^. Here, we apply a curated version of this alternative dataset to reconstruct the eToL and test the robustness of deep, inter-phyla relationships. Our results validate the monophyly of most eukaryotic supergroups, although in some cases with changes in their composition and/or internal topology, and also provide support for several novel higher-level relationships, underscoring the need to revisit the eToL with independent data.

## Results

### An independent, marker-rich, and compositionally balanced pan-eukaryotic phylogenomic dataset

To reconstruct the eToL with markers independent from the legacy datasets, we built a *de novo* phylogenomic supermatrix by mining the BUSCO Eukaryota odb9 collection, which includes 303 universal single-copy protein-coding genes^26^. For each marker, we reconstructed a phylogeny with homologs from a broad taxonomic spectrum of eukaryotic species. Several of these genes showed ancient duplications that yielded eukaryote-wide paralogs, which we further separated into independent orthologous subfamilies. We thus identified 405 orthologous markers, and excluded 128 after rigorous manual curation because they lacked a clear phylogenetic signal or exhibited non-vertical, complex evolutionary histories (see Methods and Supplementary Table 1). We finally kept 277 conserved proteins for further analyses.

The primary strength of this dataset is its independence from the current core marker set two decades ago^9^ (Supplementary Fig. 1). We evaluated the novelty of our BUSCO-based dataset by comparing it with two recently published phylogenomic marker sets (hereafter referred to as Strassert21^30^ and Tice21^3^) containing 320 and 240 markers, respectively. 210 markers were shared between Strassert21 and Tice21, accounting for ∼66% and ∼87% of these datasets, respectively. By contrast, our BUSCO-derived supermatrix showed only ∼23% overlap since 213 of our 277 markers (77%) were unique to this study (Fig. 1a). This minimal overlap provides the first opportunity to test deep eukaryotic relationships with a dataset that is largely independent from the historical consensus.

**Fig. 1.**
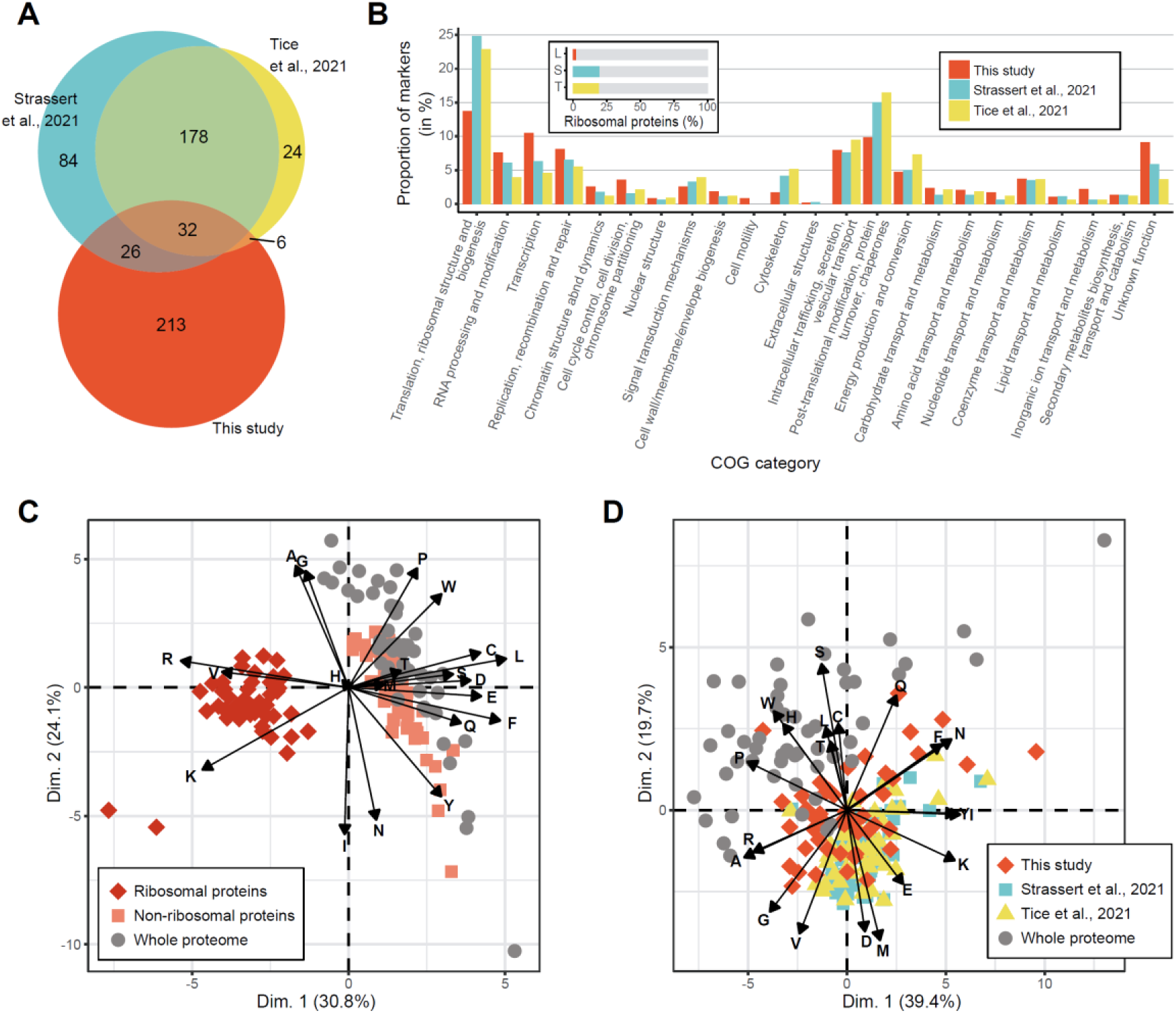
Characteristics and comparison of pan-eukaryotic phylogenomic datasets. a) Venn diagram showing the number of markers shared between, and unique to, the datasets from Strassert21, Tice21, and this study. b) COG categories represented in each phylogenomic dataset. The inset shows the proportion of ribosomal proteins in the different datasets. c) Principal component analysis (PCA) based on the amino acid composition of ribosomal proteins, non-ribosomal markers and whole proteomes for 51 diverse eukaryotic species in the dataset from this study. Similar PCAs for the Strassert21 and Tice21 datasets are provided in Supplementary Fig. 2. d) PCA based on the amino acid composition of the three phylogenomic datasets and whole proteomes for 51 diverse eukaryotic species (Supplementary Table 2).

Beyond independence, our dataset shows important shifts in the functional and compositional nature of the markers used. Legacy datasets are largely enriched in ribosomal proteins (19.3% of markers in Strassert21 and 19.1% in Tice21, see Fig. 1b). Although widely used as phylogenetic markers, ribosomal proteins—by virtue of physically interacting and co-evolving within the highly constrained structure of the ribosome—are enriched in basic amino acids (notably arginine (R) and lysine (K)) compared to the proteome average, a compositional bias known to affect phylogenetic reconstruction^31–33^. Consequently, phylogenomic datasets enriched in ribosomal proteins can skew their amino acid composition away from that of whole proteomes. Indeed, Principal Component Analysis (PCA) reveals that these R+K-rich ribosomal proteins occupy a distinct compositional space, far from the whole proteome profiles (Fig. 1c). Because legacy datasets are enriched in these markers, their overall composition is also skewed away from the proteome average. By contrast, our BUSCO-derived markers, functionally diverse and with fewer ribosomal proteins (2.1% of our dataset), have an amino acid composition that closely tracks the centroid of the whole proteomes (Fig. 1c, Supplementary Fig. 2). Therefore, our dataset minimizes this specific source of compositional bias, which may have affected previous phylogenomic reconstructions. Moreover, the limited overlap with legacy marker sets provides a critical opportunity to test the robustness and stability of the evolutionary relationships, especially the deep ones, reported in previous pan-eukaryotic phylogenies.

### Phylogenomic inference of the eToL with the new dataset

To evaluate the phylogenetic signal of our BUSCO-derived markers, we assembled a comprehensive dataset of 741 eukaryotic proteomes from the EukProt v2 database^34^. This pool was supplemented with genomic and transcriptomic data for undersampled “orphan” lineages to represent all known phyla (Supplementary Table 2). To establish a benchmark for dataset validity, we categorized these taxa into 29 uncontroversial, high-rank eukaryotic groups based on current morphology-and sequence-based consensus (Supplementary Table 2). After rigorous quality control to remove species displaying low marker coverage and obvious contamination, we retained 651 high-quality proteomes (Supplementary Table 2). In a second step, to decrease the computational burden in subsequent analyses, we curated subsamples of 264 and 61 taxa keeping a taxonomically balanced coverage of eukaryotic diversity (Supplementary Table 2).

We reconstructed a Maximum Likelihood (ML) phylogeny of the 651 taxa with the 277-marker supermatrix using the ELM+C60+G4 substitution model, which implements the recently published ELM exchangeability matrix specifically designed for pan-eukaryotic phylogenomic analyses^35^. The deep relationships among eukaryotic lineages recovered in this phylogeny (Fig. 2) were highly consistent with results from previous studies^3,4,30,36–38^. Notably, we found full statistical support for the monophyly of all previously recognized eukaryotic phyla, as well as several supergroups, such as Obazoa (Opisthokonta, Apusomonadida, Breviatea)^14^, CRuMs (Collodictyonidae, *Rigifila*, *Mantamonas*)^15^, and SAR^11^ (Fig. 2). We also recovered strong support for the deepest eukaryotic divisions including the monophyly of Podiata^39^, Opimoda^25^, and Diaphoretickes^1^. These results indicated the presence of strong vertical phylogenetic signal in our dataset.

**Fig 2.**
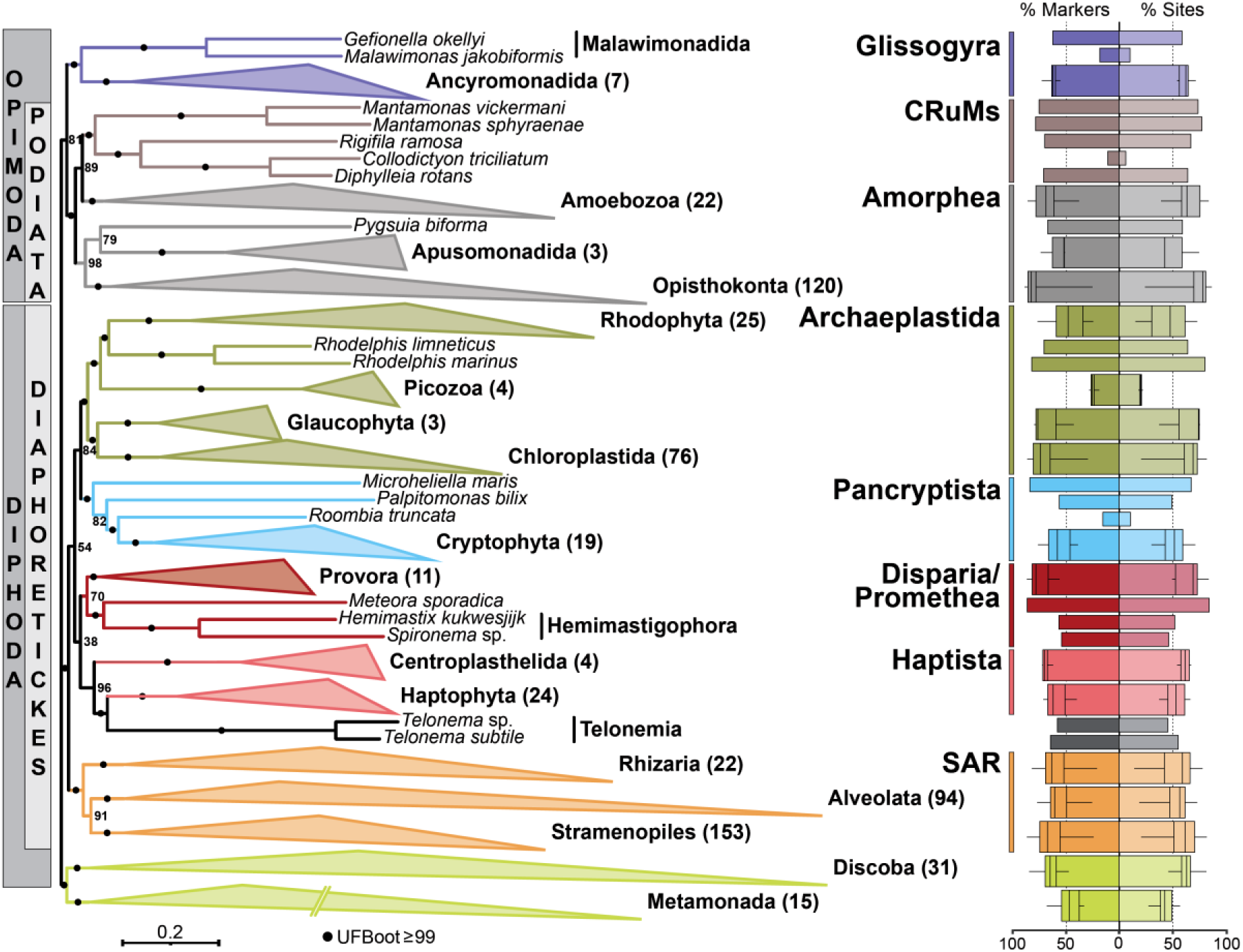
Phylogenetic tree of eukaryotes inferred from an independent phylogenomic dataset. The tree contains 651 taxa and was reconstructed using 277 markers (67,744 amino acid sites) with IQ-TREE under the ELM+C60+G4 substitution model. Support values on branches correspond to ultrafast bootstrap (UFB). Numbers next to collapsed clades indicate the number of taxa. Branch lengths for Metamonada were halved for display purposes. Bar-and boxplots on the right show marker (left) and site (right) coverages. For groups with more than two species, data are represented as box plots where the middle line is the median, the lower and upper hinges indicate the first and third quartiles, the upper whisker extends from the hinge to the largest value no further than 1.5× the interquartile range (IQR) and the lower whisker extends from the hinge to the smallest value at most 1.5× IQR.

### Robust support for most eukaryotic phyla and supergroups

To rigorously assess the stability of the relationships recovered in our first phylogeny while limiting the computational burden, we performed three targeted robustness analyses using the 264-taxon dataset. First, to mitigate the effects of mutational saturation, a known source of phylogenetic artefact^40^, we sequentially removed the fastest-evolving sites from the multiple alignment and monitored the stability of key nodes across stepwise increments of data filtering. Second, to evaluate the consistency of the phylogenetic signal and potential conflicts among gene sets, we inferred phylogenies from random marker subsamples, generating replicates using 20% and 60% of the 277 markers. Finally, to validate our results using Bayesian Inference (BI) with the sophisticated CAT+GTR+G4 substitution model^41^, which is considered more robust to Long Branch Attraction (LBA) but computationally prohibitive for large datasets, we reduced the dataset to 61 representative taxa (Supplementary Table 2). The results of these analyses concerning specific eukaryotic clades are described in the following sections.

### Opimoda and the new supergroup Glissogyra (Ancyromonadida-Malawimonadida)

In our analyses, Malawimonadida and Ancyromonadida consistently formed a well-supported monophyletic group, sister to all other Opimoda (Figs. 2, 3a). Although observed in previous analyses with variable support^3,4,37^, the robustness of this group was not analyzed in detail. We observed that it remained highly supported by our dataset even after removing more than 50% of the fastest-evolving sites, as well as in all samples of 60% markers (Figs. 3b, 3c), and in BI analysis (Supplementary Fig. 3). Interestingly, Ancyromonadida and Malawimonadida are the only opimodan lineages that possess the unique family A DNA polymerase rdxPolA, possibly representing a synapomorphy for this new clade^42^. Given the robustness of this clade across methodological treatments and independent marker sets (see below), we propose to formally name it Glissogyra (see Taxonomic description below).

**Fig 3.**
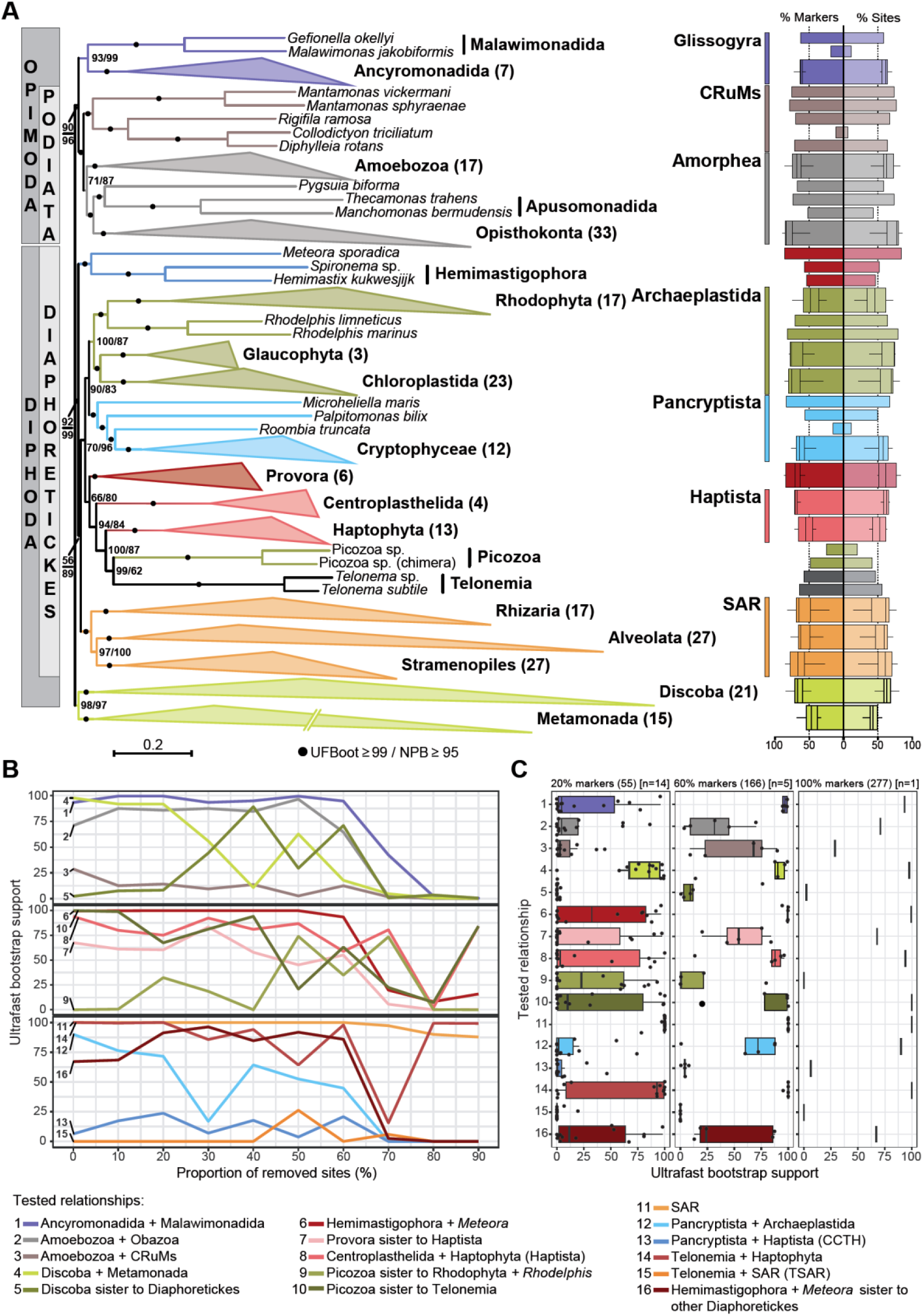
Phylogenomic analyses of deep relationships in the eToL. a) Maximum likelihood tree containing 264 taxa, reconstructed using 277 markers (66,325 amino acid sites) and the ELM+C60+G4-PMSF substitution model. Support values on branches correspond to ultrafast bootstrap (UFB) and PMSF non-parametric bootstrap (NPB), respectively. Numbers next to collapsed clades indicate the number of taxa. Branch lengths for Metamonada were halved for display purposes. Bar-and boxplots on the right show marker (left) and site (right) coverages. For groups with more than two species, data are represented as box plots where the middle line is the median, the lower and upper hinges indicate the first and third quartiles, the upper whisker extends from the hinge to the largest value no further than 1.5× the interquartile range (IQR) and the lower whisker extends from the hinge to the smallest value at most 1.5× IQR. Alternative topologies are displayed in Supplementary Fig. 3. b) Ultrafast bootstrap support for specific branches following the incremental removal of the fastest-evolving sites from the full marker concatenation. c) Ultrafast bootstrap support for specific relationships after randomly subsampling 20% (left panel), 60% (center panel) and 100% (right panel) of the 277 markers. Sample sizes were chosen based on Brown et al. (2018). Data are represented as box plots where the middle line is the median, the lower and upper hinges indicate the first and third quartiles, the upper whisker extends from the hinge to the largest value no further than 1.5× the interquartile range (IQR) and the lower whisker extends from the hinge to the smallest value at most 1.5× IQR.

While the monophyly of Obazoa (Opisthokonta, Breviatea and Apusomonadida^14^) was strongly supported in all analyses, the position of Amoebozoa shifted from sister group to CRuMs in the 651-taxa tree (Fig. 2) to the traditional position as sister group to Obazoa in the 264-taxa ML tree and in the 61-taxa BI tree (Fig. 3a; Supplementary Fig. 3). The position of Amoebozoa was also sensitive to data subsampling: removal of fast-evolving sites supported Amoebozoa as sister to Obazoa (Fig. 3b), consistent with the traditional Amorphea grouping; however, subsampling of 60% of markers alternatively supported Amoebozoa sister to CRuMs (Fig. 3c), which was also the second-best supported topology by the full 264-taxa dataset (Supplementary Fig. 4a). Therefore, the position of Amoebozoa remained uncertain and we examined it using a different set of markers (see below).

### Long-branch attraction in Excavata lineages

While Malawimonadida branched confidently with Ancyromonadida, the two other ‘excavate’ lineages, Metamonada and Discoba^1^ formed a highly-supported monophyletic group (Figs. 2, 3). Their monophyly has been proposed^20^ and debated over the years^4,5,15,17^. Indeed, Metamonada and Discoba exhibit relatively long branches compared to other eukaryotes (Figs. 2, 3), and their grouping suspected to be the result of a long-branch attraction (LBA) artifact for a long time^7^. In agreement with this view, removal of the 40% fastest-evolving sites from the alignment caused their monophyly to collapse, placing Discoba robustly within Diphoda as sister to the well-supported Diaphoretickes (Fig. 3b), consistent with previous studies^4^. In the marker subsampling analysis, the monophyly of Metamonada and Discoba remained the most supported topology, but with weaker support than with the full concatenation (Fig. 3c). This decrease in support suggested a possible systematic bias in the markers of the two lineages, which would increase the effect of LBA and/or compositional bias when increasing the amount of sequence data^43^. Interestingly, we also found Discoba as sister to Diaphoretickes when using the CAT+GTR+G4 model in BI analyses, considered to be less prone to LBA artifacts^44^.

### Telonemids and haptophytes are sister groups

In a significant deviation from a number of previous studies supporting a relationship of Telonemia with SAR (the so-called TSAR supergroup)^37,45^, Telonemia was consistently placed in our trees as sister to Haptophyta with high support (Figs. 2, 3a). In other recent analyses that did not recover the TSAR monophyly^3,4,36,46,47^, Telonemia and their sister group (either Haptophyta or Hemimastigophora) were always monophyletic with the SAR clade. However, in all of our trees, Telonemia+Haptophyta and SAR appeared to be distantly related (e.g., Figs. 2, 3a). Furthermore, the Telonemia+Haptophyta relationship remained fully supported after removing any number of fast-evolving sites (Fig. 3b) and was also the best supported topology in the marker subsampling analysis (Fig. 3c).

### Resolution of enigmatic orphan lineages

Our analyses also clarified the position of several other orphan lineages (i.e., those lacking clear phylogenetic affinities, often with few available sequenced genomes or transcriptomes) and highlighted instances where compositional bias may drive the tree topology.

In agreement with recent studies^4,36–38^, we obtained full support for the monophyly of Hemimastigophora and *Meteora sporadica* (Figs. 2, 3a), regardless of the number of fast-evolving sites removed (Fig. 3b). These two lineages were placed with moderate support at the base of the Diaphoretickes. Our 264-taxa analysis placed Provora as sister to Haptista, similarly to previous analyses^23^, but in contrast with our 651-taxa tree (Fig. 2) and other recent studies that placed them as sister to Hemimastigophora and *M. sporadica*^4,36–38^. While the latter placement was recovered in only 1% of our non-parametric bootstrap (NPB) trees (Supplementary Fig. 4b), the Provora-Haptista relationship remained the most supported one when removing the fastest-evolving sites and when subsampling markers (Figs. 3b, 3c).

Picozoa have been proposed to be sister to Rhodophyta and *Rhodelphis* within the supergroup Archaeplastida^2^, a placement confirmed by our 651-taxa ML tree (Fig. 2) and our BI phylogeny (Supplementary Fig. 3). Intriguingly, this clade was placed as sister to Telonemia and Haptophyta with moderate to high support in the 264-taxa ML tree (Fig. 3a), whereas the alternative position within Archaeplastida was recovered in only 10% of the NPB trees (Supplementary Fig. 4b). However, we observed that Picozoa and Telonemia were grouped in an amino acid composition-based hierarchical clustering of the 264 taxa (Supplementary Fig. 5), suggesting that compositional bias drove the placement of Picozoa out of Archaeplastida in this particular ML inference. Supporting this hypothesis, the reconstruction of a phylogeny without Telonemia recovered with full support the expected position of Picozoa within Archaeplastida (Supplementary Fig. 6). These results suggest that the phylogenetic placement of Picozoa is sensitive to compositional bias and to similarities in amino acid composition to Telonemia.

Finally, the monophyly of the supergroups Archaeplastida, Pancryptista and Haptista (plus Telonemia and Provora) appeared reasonably supported (96% NPB in the 264-taxa tree, but only 54% UFB in the 651-taxa tree) (Figs. 2 and 3a). Although the relationships within this clade remained unclear, its monophyly has only been recovered a few times before, most recent phylogenomic studies instead supporting a sister relationship between Haptista and SAR^4,36–38,47^. Moreover, the moderate support for the monophyly of Archaeplastida and Pancryptista contrasts with other recent studies^3,30^, pointing to a potentially incongruent signal with previous phylogenomic markers.

### Incongruent phylogenetic signal between marker sets

The 64 markers shared between our dataset and the Strassert21 and/or Tice21 datasets (Fig. 1a) accounted for a substantial fraction of our original supermatrix (∼20,000 sites) and may have masked signal from markers unique to our dataset. Therefore, to disentangle the effects of marker choice and taxon sampling, we compared ML phylogenies inferred from our dataset-specific markers with those based on previously published sets. After updating the Strassert21 and Tice21 datasets to match our sampling of 264 taxa, we generated two non-overlapping concatenations containing, respectively, 213 markers unique to this study (Leroy-specific (LS)) and 350 markers from the combined Strassert21+Tice21 datasets (ST).

The overall topologies of the trees reconstructed from the LS and ST concatenations were similar. In fact, although the resulting supermatrices differed markedly in length (47,217 sites for LS vs. 94,293 for ST), which could influence support as longer supermatrices tend to yield higher bootstrap values^48^, both supported most well-recognized eukaryotic phyla and supergroups (Fig. 4). Notably, the monophyly of Glissogyra (Ancyromonadida + Malawimonadida) was strongly supported in both trees, as were classical supergroups such as SAR and Archaeplastida. However, the two tree topologies also showed several significant conflicts.

**Fig 4.**
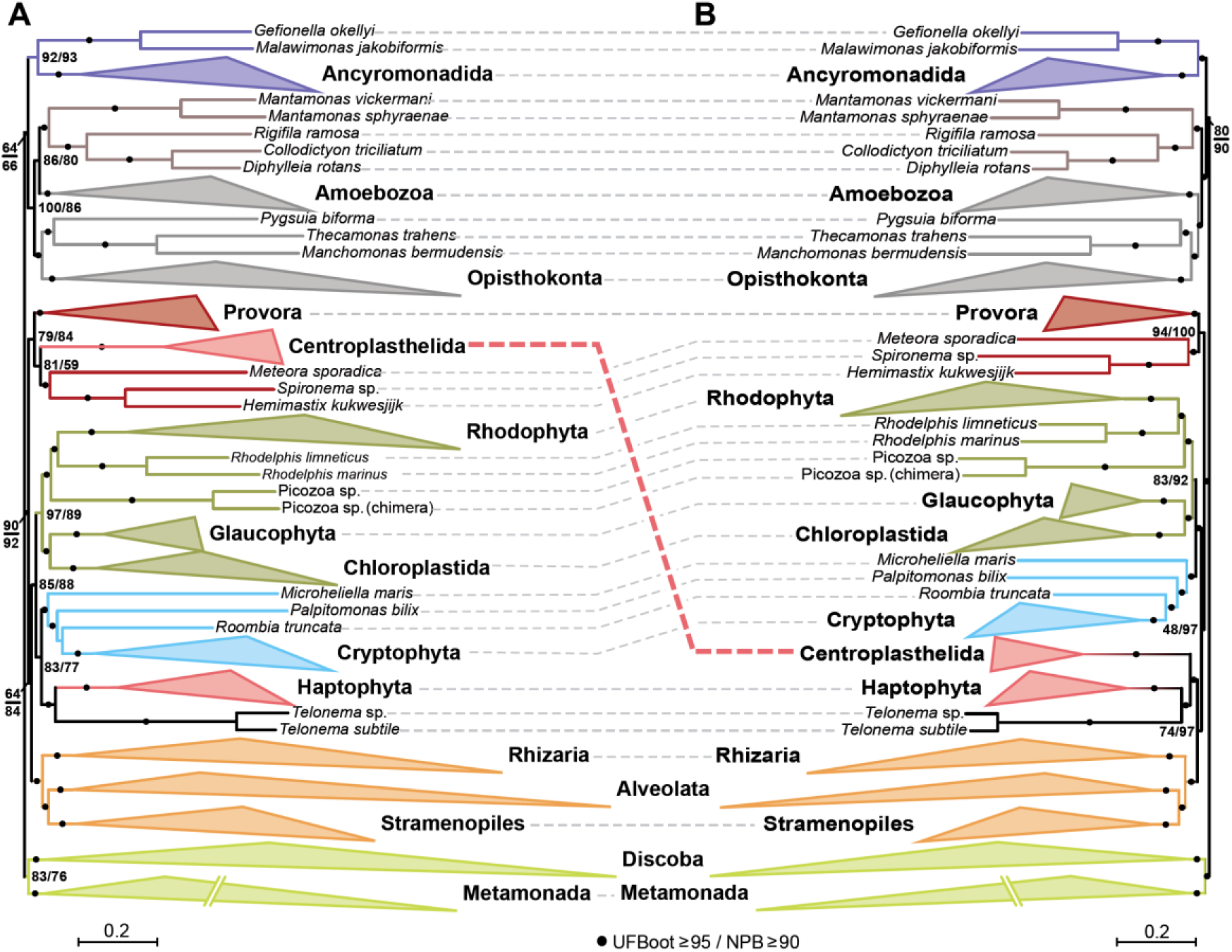
Comparison of ML phylogenetic trees based on mutually-exclusive phylogenomic datasets. a) tree based on the Leroy-specific (LS) concatenation (213 markers, 47,217 sites). b) tree based on the Strassert-Tice (ST) concatenation (350 markers, 94,293 sites). Support values on branches correspond to ultrafast bootstrap (UFB) and PMSF non-parametric bootstrap (NPB). Branch lengths for Metamonada were halved for display purposes. The branch of Centroplastelida showed the most dramatic shift in position between the two trees, as indicated by the dashed red line.

Within the Opimoda, while the ST tree recovered Amoebozoa and Obazoa as sister groups (forming the Amorphea), the LS tree placed Amoebozoa as sister to CRuMs with moderate support. This indicates a divergent phylogenetic signal for this group between the two datasets, consistent with our previous finding that the Amoebozoa + CRuMs topology was the second most supported when using the full 277-marker dataset (Supplementary Fig. 4a).

However, most discordances appeared in the Diaphoretickes region of the tree (Fig. 4). The LS tree placed Centroplasthelida within the Provora-Hemimastigophora-*Meteora* clade, rendering Haptista polyphyletic, differing from the ST phylogeny (Fig. 2, 4). This incongruence is particularly interesting because Centroplasthelida and Haptophyta share filiform microtubule-supported feeding structures^49^. However, similar feeding structures occur in distantly related groups (e.g., Radiolaria^50^), which, combined with our phylogenomic analyses, suggest that these features may be convergent rather than synapomorphic.

Along with the breakup of Haptista, the LS phylogeny grouped Pancryptista with Haptophyta and Telonemia rather than with Archaeplastida, in contrast with the 651-taxa tree (Fig. 2) and the ST phylogeny (Fig. 4). This assemblage is reminiscent of the CCTH hypothesis (i.e., the grouping of Cryptophyta, Centroplasthelida, Telonemia, and Haptophyta^12^). Related to this, while the LS tree recovered the deep-branching placement of SAR within Diaphoretickes observed in our previous analyses (Fig. 2), the ST phylogeny placed them as sister to Haptophyta+Telonemia+Centroplasthelida (Fig. 4), as seen in previous studies^4,12,21,36–38^. These disagreements between different datasets indicate that the phylogenetic relationships within the Diaphoretickes cannot be considered as settled, which likely stems from the rapid evolutionary radiation of these lineages that made them particularly sensitive to marker choice^49,51^. This is consistent with the observation that, to the exception of the well-supported deep-branching position of Hemimastigophora + *Meteora*, this region of the tree also remained unresolved in our BI tree (Supplementary Fig. 3).

The monophyly of Discoba and Metamonada, although recovered in both LS and ST phylogenies, was less supported in the LS tree. This result was also consistent with our fast site-removal analysis (Fig. 3b), reinforcing the idea that the monophyly of Discoba and Metamonada observed in many studies is most likely due to an LBA artefact^43^. This was supported by our BI analysis using the CAT+GTR+G4 model, which explicitly accounts for site-specific compositional heterogeneity and is considered more robust to LBA (Supplementary Fig. 3). In this analysis, Discoba branched within Diphoda as sister to Diaphoretickes (with posterior probability = 0.94), consistent with the recent studies proposal of a major split of eukaryotes between the two groups ‘Opimoda+’ (including Metamonada) and ‘Diphoda+’ (including Discoba)^5^.

Finally, we concatenated all the 563 markers analyzed in this study (213 LS + 350 ST ones), generating an alignment of 141,510 amino acid positions. Unsurprisingly, due to the large proportion of sites coming from the ST markers (∼67%), the resulting tree was effectively identical to the ST phylogeny (Fig. 4b and Supplementary Fig. 7). Notably, we recovered full statistical support for the monophyly of Ancyromonadida + Malawimonadida, as well as for the monophyly of Telonemia + Haptophyta.

## Discussion

Over the past two decades, in parallel with methodological advances in phylogenetic reconstruction, the eToL has been refined largely by adding more taxa to the same core set of protein markers. Despite the substantial progress achieved through this approach, it remained necessary to build an alternative dataset of independent markers to (i) test the robustness of previous findings and (ii) potentially provide additional signal to resolve uncertain branches. To this end, we built a largely non-overlapping BUSCO-derived marker set. The resulting phylogenies corroborate the monophyly of most well-established high-rank groups and recover robust support also for a number of deep divisions, indicating that substantial vertical signal is retained in this dataset. At the same time, our analyses strengthen support for relationships that were previously only moderately supported (most notably a basal opimodan clade uniting Ancyromonadida and Malawimonadida) and identify regions of the tree that remain sensitive to marker choice and model assumptions, including the internal branches of Diaphoretickes, the position of Amoebozoa, and the apparent monophyly of ‘Excavata’ lineages (Discoba and Metamonada).

The recovery in some analyses of a CCTH-like clade (i.e., Cryptophyta, Centroplasthelida, Telonemia, and Haptophyta) has implications for hypotheses concerning the evolution of photosynthesis in eukaryotes, particularly in algae containing red-algal-derived plastids. Under several serial endosymbiosis models, the plastid donor for haptophytes was proposed to be an ancestral cryptophyte^30^, consistent with phylogenetic analyses of plastid data^52^ and of the SELMA translocon proteins^53^. Our nuclear-marker phylogenies support the alternative possibility that cryptophytes and haptophytes may share a plastid-endowed ancestor, which would shift the explanatory burden to subsequent plastid losses in close relatives of these lineages. Nevertheless, although plastid loss is increasingly documented^54^, this remains a challenging scenario.

More broadly, the overall agreement between the BUSCO-derived and legacy marker sets in supporting most eukaryotic phyla and supergroups strengthens the validity of the few-supergroup structure of eukaryotic phylogeny^1^ and suggests that the reconstruction of a consensus eToL is an attainable goal (Fig. 5). Improved genomic sampling of poorly known lineages, continued development of eukaryote-adapted mixture substitution models, and understanding the causes of incongruence between conflicting gene phylogenies will be essential to distinguish historical signal from artefacts and to converge on a stable, well-supported eukaryotic tree.

**Fig. 5.**
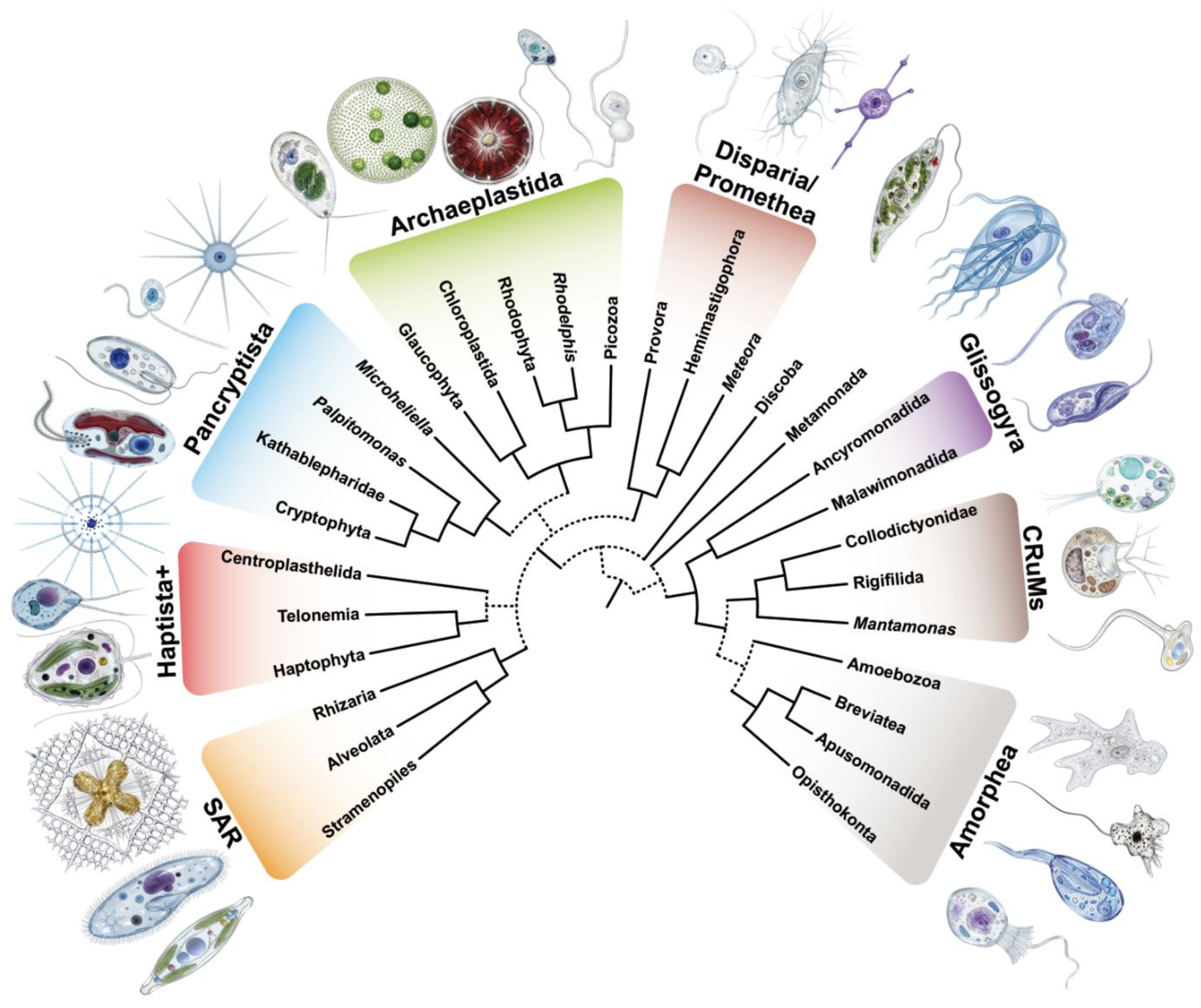
Consensus eukaryotic Tree of Life. This tree, based on this work and other recent phylogenomic analyses, shows the major eukaryotic supergroups. Consensual branches are shown as solid lines, whereas relationship not yet fully resolved are indicated with dashed lines. The supergroup ‘Haptista+’ refers to the classical Haptista (Haptohyta + Centroplasthelida) plus Telonemia.

**Taxonomic description Glissogyra** Leroy et al. 2026

The least inclusive clade of eukaryotes containing *Ancyromonas sigmoides* Saville-Kent 1880^55^ and *Malawimonas jakobiformis* O’Kelly and Nerad 1999^56^. This is a node-based definition in which all of the specifiers are extant; it is intended to apply to a crown clade.

Etymology: From Greek glíssa (γλῖσσα), sticky substance or something that adheres, and gyros (γυρος), a circle or ring, referring to the characteristic gliding and swimming motility patterns observed in representatives of both groups.

## Methods

### Taxon sampling and construction of proteome datasets

We collected all available proteomes from the EukProt v2 database^34^, to which we added missing taxa with important phylogenetic positions from the EukProt v3 database. In addition, poorly represented lineages in EukProt (i.e., fewer than 5 proteomes) with key phylogenetic positions were supplemented with published data from other databases^23,46,57–61^. When only transcriptome sequences were available, open reading frames were predicted with TransDecoder v5.7.0 (Haas, https://github.com/TransDecoder/TransDecoder) with default parameters (except the minimal peptide size set at 50), followed by dereplication with CD-HIT v4.8.1^62^ at 100% identity to remove redundant peptides.

Prokaryotic contaminations were identified in and removed from all the 770 collected proteomes (see Supplementary Table 2) by searching against the EggNOG v6.0 database^63^ using DIAMOND v2.0.11.149^64^ (parameters: blastp-e 1e-10-k 20). Sequences that had ≥50% prokaryotic hits were considered prokaryotic and removed; proteomes in which ≥25% of sequences met this criterion were excluded entirely.

Proteome completeness was then assessed using the BUSCO v3 dataset^26^ (OrthoDB v9). We used the hmmsearch tool from the HMMer v3.3.2 toolkit^65^ (--domE 1e-5) to identify up to the 3 best hits of each BUSCO marker in each proteome. We only kept proteomes with ≥30% of markers identified (≥20% for poorly represented lineages). This filtering step yielded a final set of 675 taxa.

### Identification of new phylogenomic markers

For each of the 303 BUSCO v3 markers^26^, we grouped the hits of each proteome into single ‘per-marker group’ files. Each file was aligned using MAFFT FFT-NS-2 v7.310^66^ (default parameters), trimmed using BMGE v1.12^67^ (parameters:-g 0.5-h 0.5-b 3), and used to infer a maximum likelihood (ML) phylogenetic tree using IQ-TREE v2.0.3^68^ under the LG+G4 substitution model. Marker groups in which ≥20% of their taxa had ≥2 hits were considered ancestrally duplicated, and were manually split into orthologous groups. Only orthologous groups represented by ≥25% of all 675 taxa were retained as candidate phylogenetic markers.

405 candidate markers identified in this way were then curated to keep the best representative sequence (i.e., the true orthologue) from each proteome and remove non-orthologous sequences. First, sequences from the same proteome that placed as sister in the tree were automatically pruned using ETE3^69^ to keep only the highest-scoring hit from the HMM search. Second, each marker was manually curated according to the following criteria: (i) sequences that branch outside their expected lineage, while others sequences from the same taxon do not, were removed; (ii) when no sequence from a proteome places inside its expected lineage, the highest scoring hit was kept; (iii) when multiple sequences from the same proteome placed inside their expected lineage, the highest scoring hit was kept; (iv) when multiple sequences from the same proteome placed inside their expected lineage but in different monophyletic clades, the hit branching in the clade that best matched the consensual intra-lineage phylogeny was kept. A final third round of manual curation was done to remove paralogous sequences, prokaryotic and eukaryotic contaminations, and potential horizontally transferred sequences. During this last round, we filtered out 24 proteomes that showed clear signs of abundant contamination or extreme divergence, as well as 128 markers that showed signs of orthology detection issues or that did not recover consensual monophyletic groups (Supplementary Table 1). The final dataset comprised 277 phylogenomic markers spanning 651 proteomes.

### Comparison of phylogenomic datasets

To assess the novelty and composition of our dataset relative to previously published ones, we performed comparisons with the Strassert21^30^ and Tice21^3^ datasets. We first looked for shared markers across datasets using BLAST searches. To do so, we selected 51 representative species present in all three datasets (Supplementary Table 2). For each marker, we performed pairwise BLASTp v2.12.0^70^ searches against markers from the other datasets. Hits with 100% identity and ≥99% coverage were considered matches; markers were considered identical if at least 10% of the representative species showed a match.

Second, we compared the functional categories of the markers in the different datasets by running EggNOG-mapper v2.1.10^71^ searches (parameter:-m mmseqs) against the EggNOG5 database^72^. For these searches, 10 representative species were manually selected for their extensive annotation level and presence in the EggNOG5 database (Supplementary Table 2). We manually inspected the results for each marker to identify its function and functional category. BUSCO-derived markers were further validated by BLASTp searches against the NR database (https://www.ncbi.nlm.nih.gov/refseq/about/nonredundantproteins/; web version of August 2023).

Finally, we analyzed the amino acid composition of the different phylogenomic datasets using an in-house Python script. Using the 51 taxa previously selected, we computed for each taxon the frequency of all amino acids in each of the three datasets as well as in its whole proteome.

### Species tree reconstruction

We generated two initial ML trees: one using all 651 proteomes and one using a curated subset of 264 manually selected proteomes representing a balanced and broad sampling of all major eukaryotic lineages (Supplementary Table 2). In both cases, individual markers were aligned using MAFFT L-INS-I v7.310^66^ (parameter: --bl 30), trimmed with BMGE v1.12^67^ (parameters:-m BLOSUM30-g 0.2-h 0.4-b 1) and concatenated using an in-house Python script, yielding matrices of 67,744 and 66,325 sites, respectively.

For each taxon sampling, we inferred a species tree from the supermatrix using IQ-TREE v2.0.3^68^ under the ELM+C60+G4 substitution model^35^. To reduce computational time, we constrained the tree search using a backbone phylogeny constructed as follows: For each major eukaryotic lineage comprising at least 4 proteomes, we selected the 100 (out of 277) markers with the broadest taxon coverage, aligned them using MAFFT L-INS-I v7.310 (--bl 45), trimmed using BMGE v1.12 (-m BLOSUM45-g 0.2-h 0.4-b 1), and inferred a lineage-specific ML tree with IQ-TREE v2.0.3 under the ELM+C60+G4 substitution model. Each resulting tree was manually inspected for congruence with published phylogenies, and rooted according to the consensus position reported therein (Supplementary Table 3). Using an in-house Python script, we assembled a multifurcating constraint tree by collapsing all branches with low support (SH-aLRT<80 and UFB<95), leaving them free to be resolved during the full tree search.

For these large tree inferences, the substitution model sophistication comes at the expense of the computational resources needed for its use, which prevents assessing standard non-parametric bootstrap (NPB) support. To alleviate this issue, we adopted a two-step approach for the 264-taxa supermatrix. We first computed posterior mean site frequencies^73^ (PMSF) using the previously-inferred tree as a guide tree, then re-inferred the phylogeny under the ELM+C60+G4-PMSF substitution model and 500 NBP replicates.

### Phylogenomic analyses

Various analyses were conducted on the 264-taxon and 277 BUSCO-based markers supermatrix (66,325 sites).

First, to look for potential phylogeny reconstruction artifacts due to mutational saturation in fast-evolving sites, we incrementally removed them from the initial supermatrix. We ranked each site based on its evolutionary rate computed using Dist_Est v1.1.1 (http://www.mathstat.dal.ca/~tsusko/src/dist_estv1.1.1.tar.gz) and then generated alignments from the initial supermatrix by removing the fastest-evolving sites in 10% increments, from 10% to 90% of the sites. We reconstructed a tree for each alignment using IQ-TREE v2.0.3 under the ELM+C60+G4 model and the tree constraints described above. Using an in-house Python script, we tracked the ultrafast bootstrap (UFB) support for key relationships.

Second, to evaluate the robustness of the phylogenetic signal, we inferred phylogenies from random subsets of markers. Following previous studies^15^, we randomly sampled 14 replicates of 20% of all markers (i.e., 55 out of 277), and 5 replicates of 60% (i.e., 166 out of 277). For each replicate, we inferred a phylogeny using IQTREE v2.0.3 under the ELM+C60+G4 substitution model and the tree constraints mentioned above, and tracked the UFB support for key relationships.

Finally, we performed a Bayesian inference (BI) analysis on a reduced set of 61 manually selected taxa to reduce computational time. Markers were aligned, trimmed and concatenated following the same procedure used for ML analyses. Phylogenetic reconstruction was carried out using PhyloBayes-MPI v1.8c^74^ under the CAT+GTR+G4 substitution model. For this analysis, we ran 2 chains for 20,000 generations, and generated a consensus tree from these chains after a burn-in of 5,000.

### Recruitment of published phylogenomic markers

To further test the congruence between our dataset and previously-published ones, we collected the 286 markers found to be unique to the Strassert21^30^ and Tice21^3^ datasets. For each of these 286 markers, we use all sequences corresponding to taxa present in our 264-species sampling to generate HMM profiles for each marker, which were then used to fetch the 3 best hits per missing taxon using HMMer v3.3.2 (hmmer.org)^65^. The subsequent marker curation followed the same procedure as described above.

We then inferred two phylogenies based on different marker sets: i) the 213 markers unique to this study (LS, Leroy-specific), and ii) the 350 markers present in Strassert21 and Tice21. Because the ELM model is computationally intensive for large datasets, we employed a two-step approach to also obtain bootstrap support values: first, we inferred ML trees with ELM+C60+G4 to establish guide topologies. Then we estimated posterior mean site frequencies (PMSF) from these trees and re-inferred phylogenies using the faster ELM+C60+G4-PMSF model with 500 non-parametric bootstrap replicates for each dataset. This strategy preserves the phylogenetic signal of the more sophisticated model while enabling rapid bootstrap analysis.

## Acknowledgments

D.M., P.L.-G., and L.E. were supported by grants from the European Research Council (ERC Advanced grants 787904 and 101141745, and ERC Starting grant 803151, respectively). This work was also supported by the Moore Foundation GBMF9739 (to P.L.-G.) and ANR DArchFolds ANR-22-CE02-0012-02 (to D.M., P.L.-G. and L.E.). We thank P. Deschamps for help in managing our bioinformatic cluster and the Institut Français de Bioinformatique (IFB) for access to computational resources.

---

**Supplementary Fig. 1.**
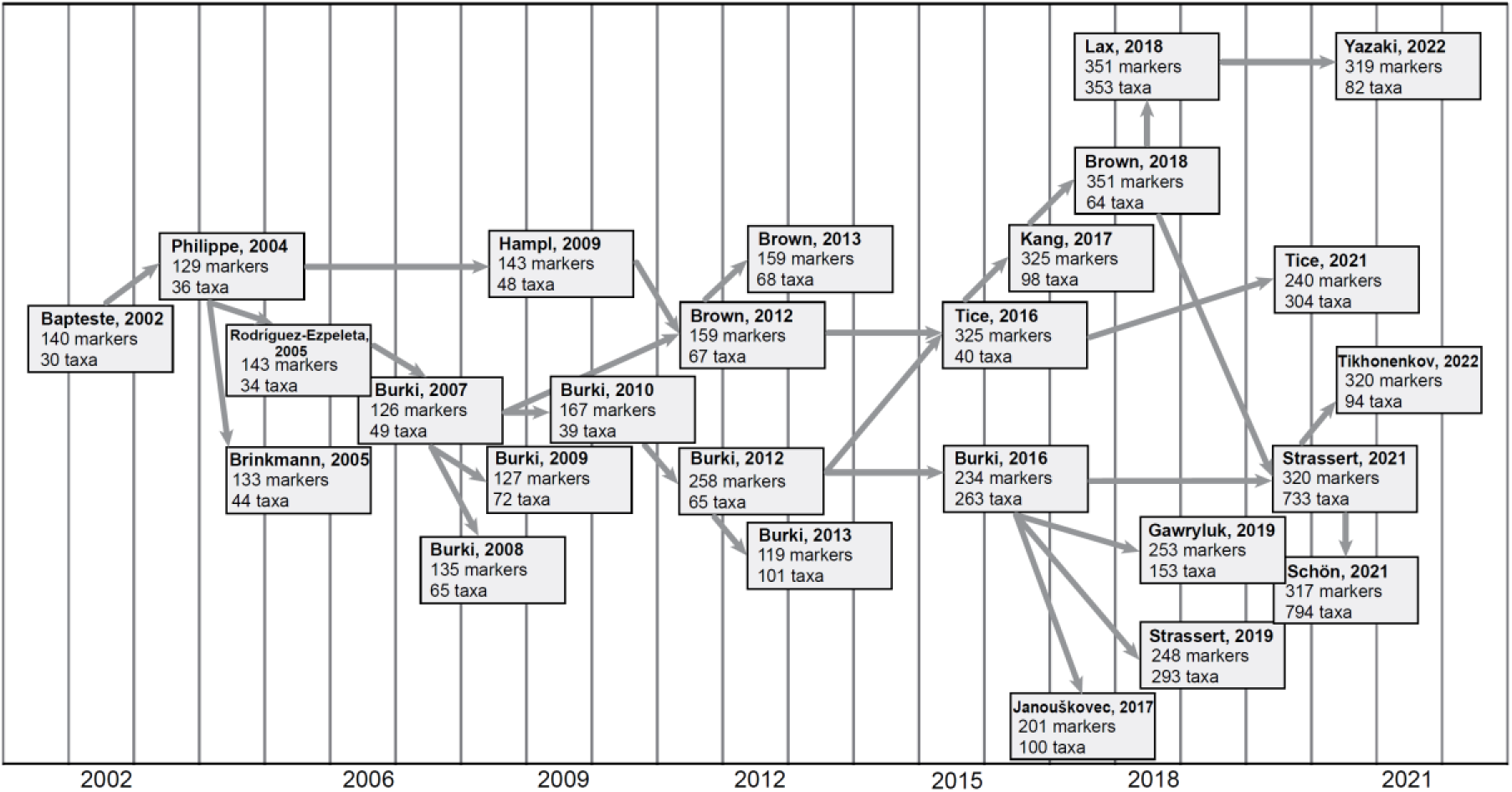
Non-exhaustive genealogy of phylogenomic datasets used for the reconstruction of the eToL. For each dataset, the number of protein markers and taxa is indicated.

**Supplementary Fig. 2.**
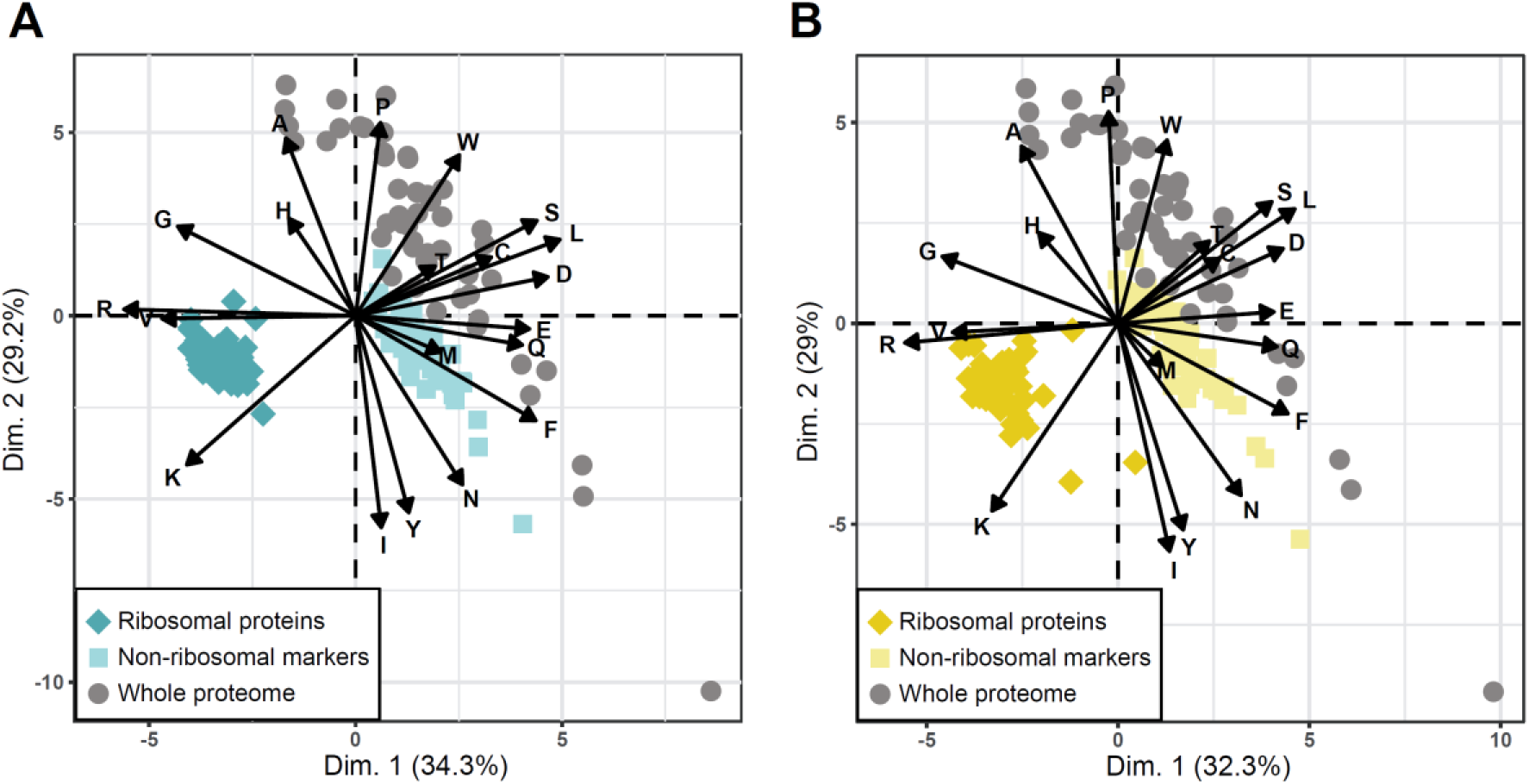
Amino acid composition of phylogenomic markers. a) Principal component analysis (PCA) based on the amino acid composition of ribosomal proteins, non-ribosomal markers and whole proteomes for 51 eukaryotes in the Strassert21 dataset^30^. b) PCA based on the amino acid composition of ribosomal proteins, non-ribosomal markers and whole proteomes for 51 eukaryotes in the Tice21 dataset^3^.

**Supplementary Fig. 3.**
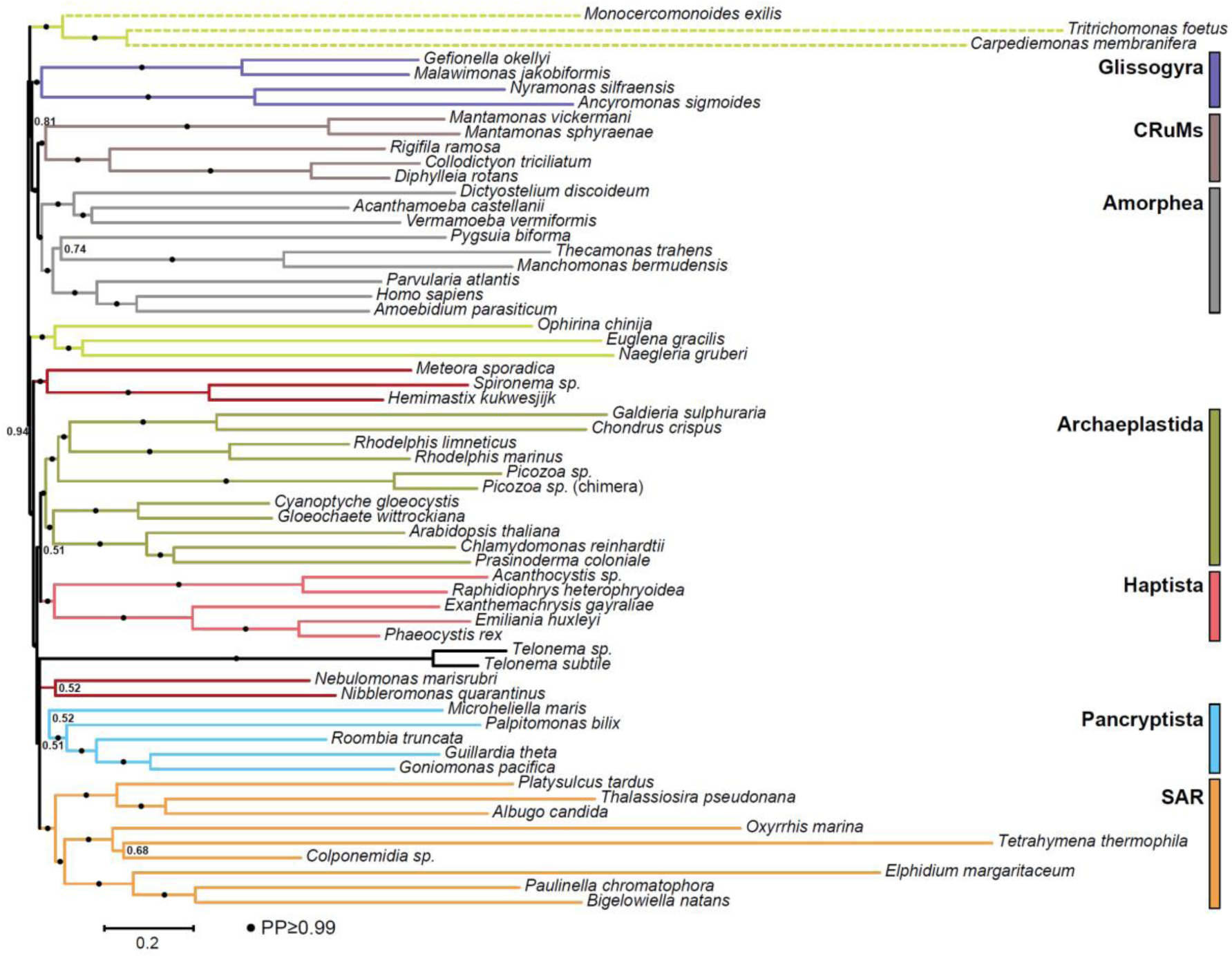
Bayesian Inference phylogenetic tree of eukaryotes. The tree contains 61 taxa and was reconstructed using 277 markers (75,546 amino acid sites) with PhyloBayes under the CAT+GTR+G4 substitution model. Support values on branches correspond to posterior probabilities (PP), estimated from a consensus of two chains. Branch lengths for Metamonada are halved for display purposes.

**Supplementary Fig. 4.**
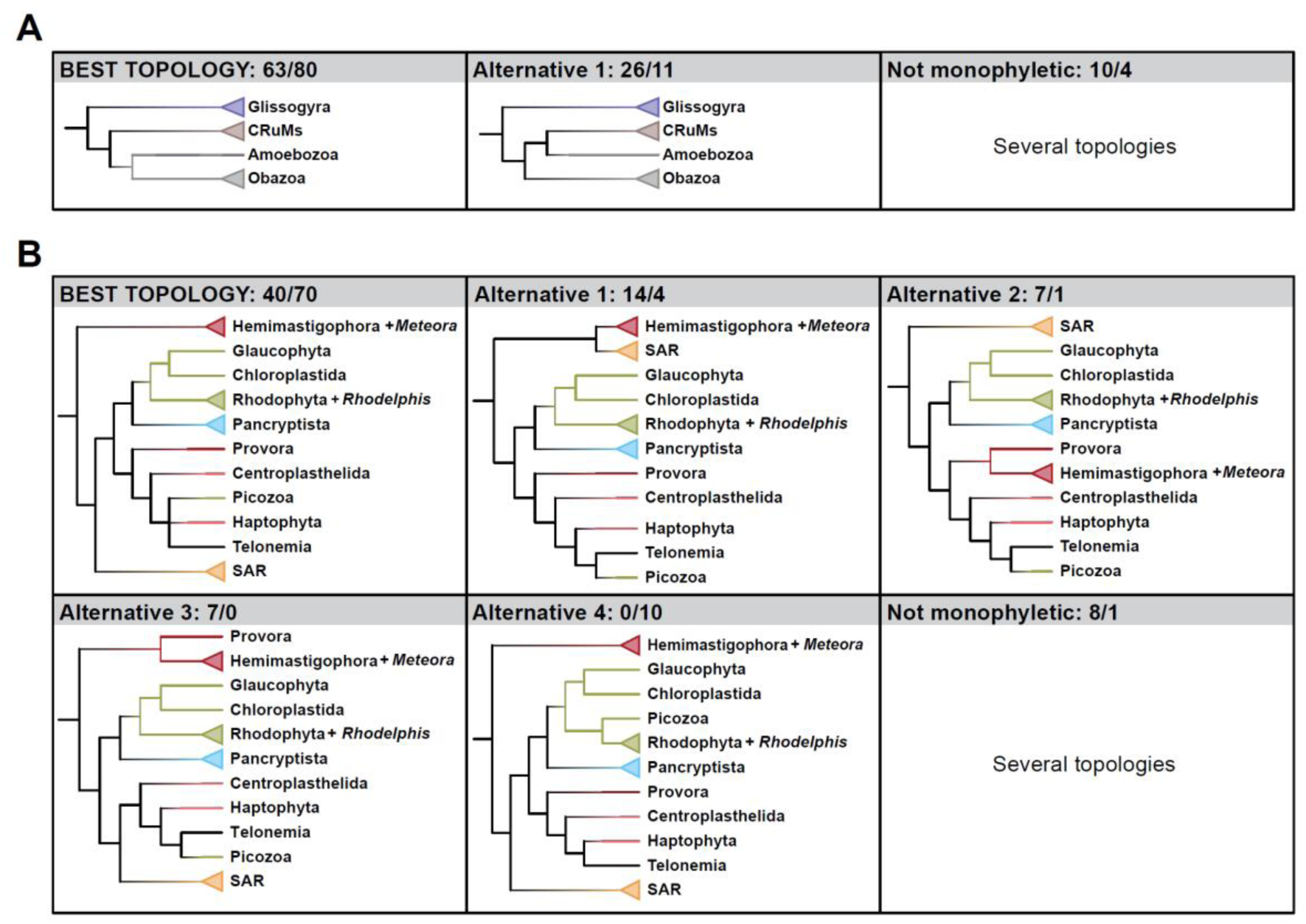
Different topologies recovered in the bootstrap replicates. a) Topologies in the Opimoda section of the tree. b) Topologies in the Diaphoretickes section of the tree. Values associated to each topology correspond to their ultrafast (UFB) and non-parametric (NPB) bootstrap support (i.e., the proportion of bootstrap replicates showing a particular topology), respectively. Only cases with ≥5% UFB or NPB support are displayed.

**Supplementary Fig. 5.**
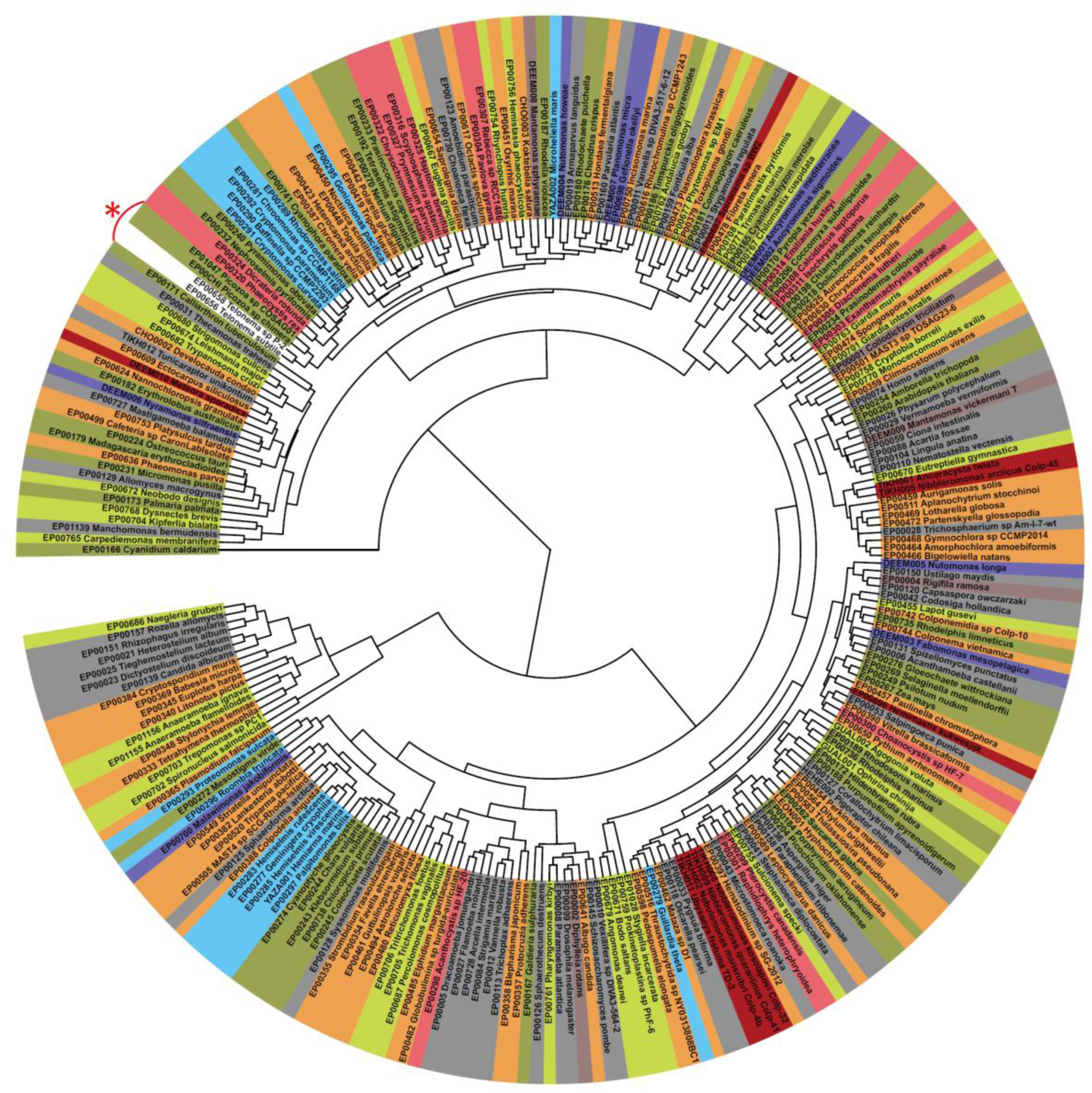
Hierarchical clustering of 264 taxa based on amino acid composition. The proportion of each amino acid was inferred based on the non-gap positions of the 66,325-sites concatenation. All 20 amino acids were used to compute Euclidian distances between each pair of taxa. Colors indicate different major taxonomic groups. The red asterisk indicates the position of Telonemia and Picozoa.

**Supplementary Fig. 6.**
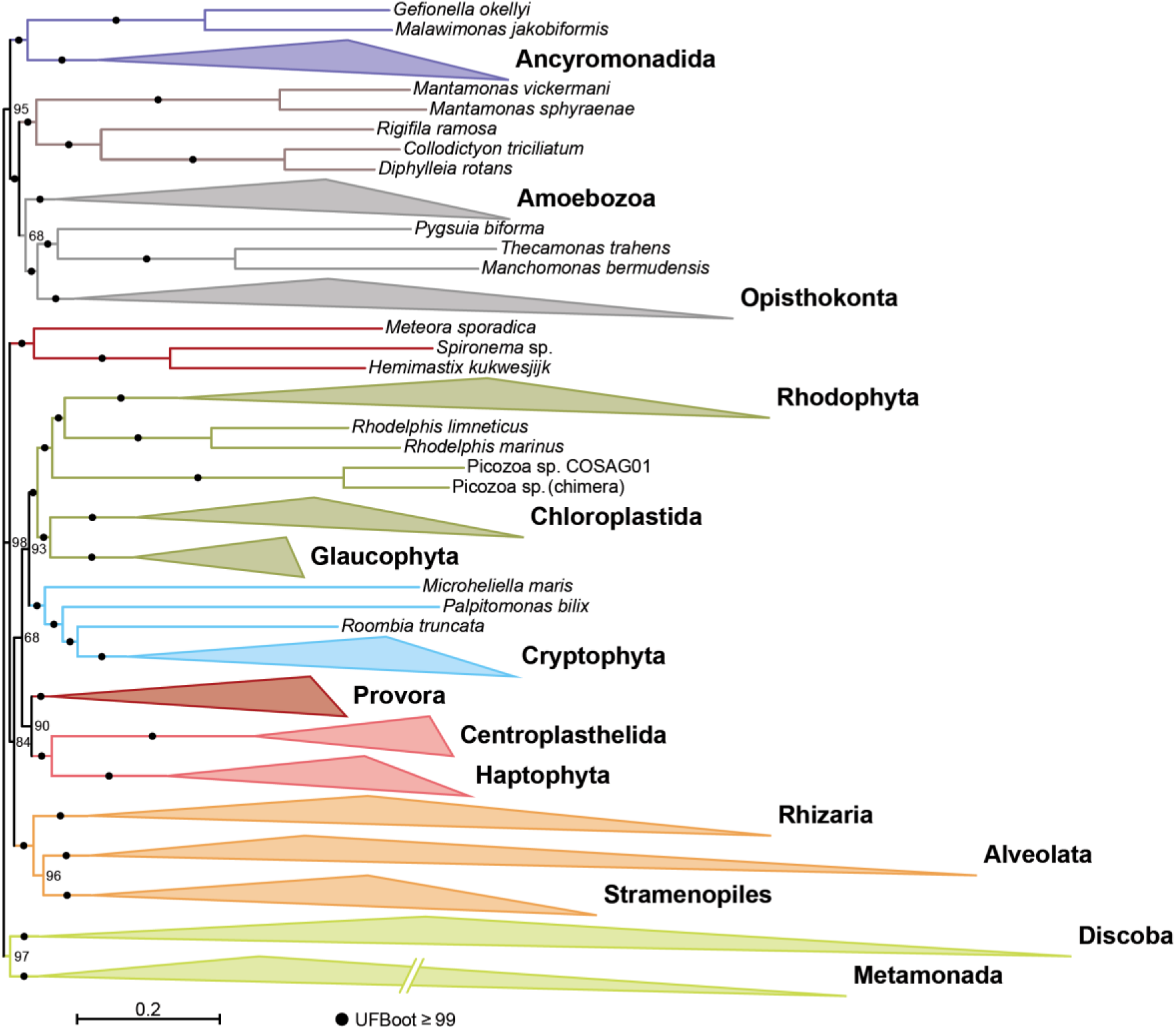
Phylogenetic tree of eukaryotes excluding telonemids. The tree contains 262 taxa and was reconstructed using 277 markers (66,325 amino acid sites) with IQ-TREE under the ELM+C60+G4 substitution model. Support values on branches correspond to ultrafast bootstrap (UFB). Branch lengths for Metamonada were halved for display purposes.

**Supplementary Fig. 7.**
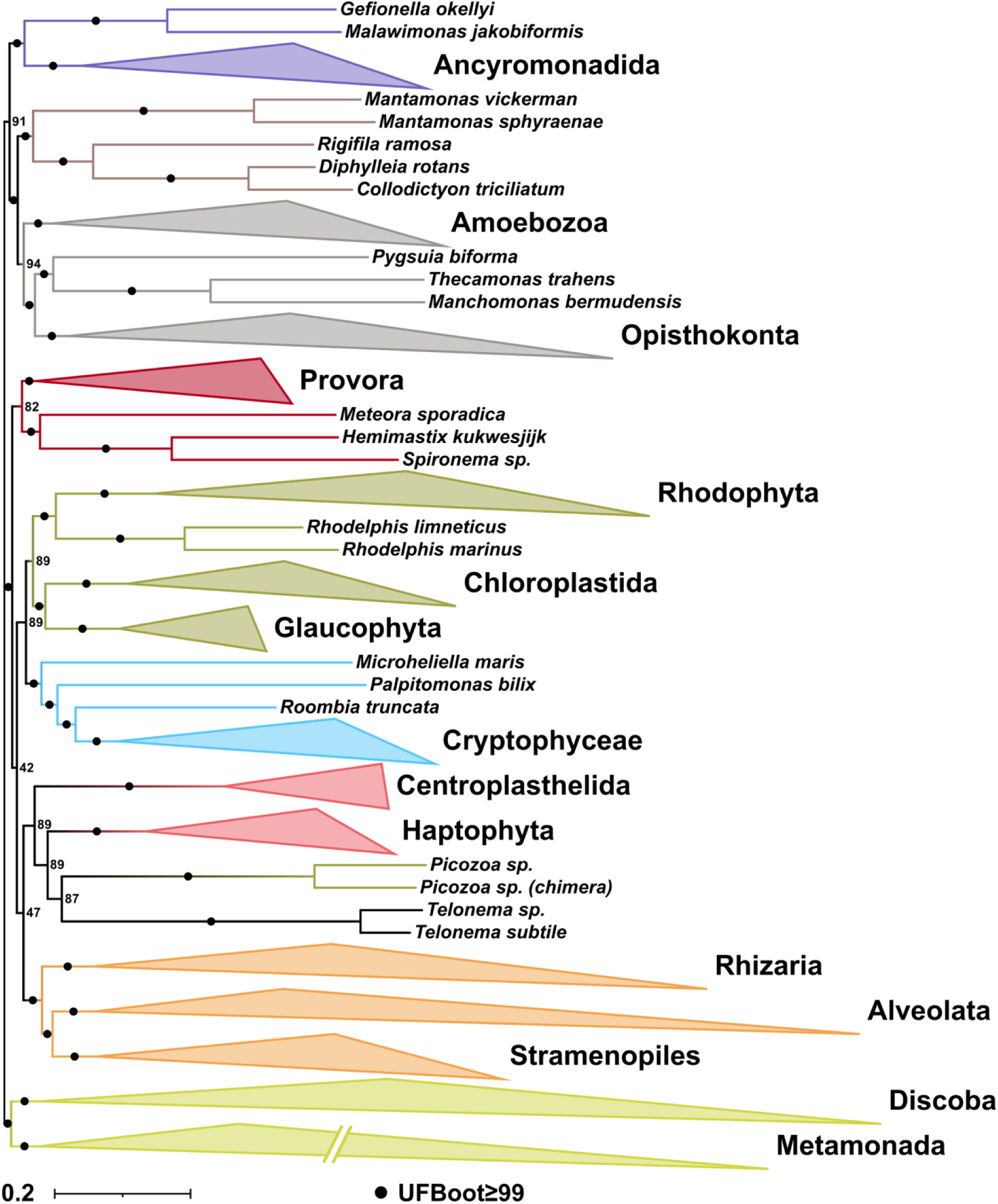
Phylogenetic tree of eukaryotes based on all available markers. The tree contains 264 taxa and was reconstructed using 563 markers (141,510 amino acid sites) with IQ-TREE under the ELM+C60+G4 substitution model. Support values on branches correspond to ultrafast bootstrap (UFB). Branch lengths for Metamonada were halved for display purposes.

## References

1. Adl, S.M. et al. The revised classification of eukaryotes. J Eukaryot Microbiol 59, 429–493 (2012).

2. Schön, M.E. et al. Single cell genomics reveals plastid-lacking Picozoa are close relatives of red algae. Nat Commun 12, 6651 (2021).

3. Tice, A.K. et al. PhyloFisher: A phylogenomic package for resolving eukaryotic relationships. PLoS Biol 19, e3001365 (2021).

4. Torruella, G., Galindo, L.J., Moreira, D. & López-García, P. Phylogenomics of neglected flagellated protists supports a revised eukaryotic tree of life. Curr Biol 35, 198–207.e4 (2025).

5. Williamson, K. et al. A robustly rooted tree of eukaryotes reveals their excavate ancestry. Nature 640, 974–981 (2025).

6. Richards, T.A. et al. Reconstructing the last common ancestor of all eukaryotes. PLoS Biol 22, e3002917 (2024).

7. Philippe, H. et al. Early-branching or fast-evolving eukaryotes? An answer based on slowly evolving positions. Proc R Soc Lond B Biol Sci 267, 1213–1221 (2000).

8. Sibbald, S.J. & Archibald, J.M. More protist genomes needed. Nat Ecol Evol 1, 145 (2017).

9. Bapteste, E. et al. The analysis of 100 genes supports the grouping of three highly divergent amoebae: *Dictyostelium*, *Entamoeba*, and *Mastigamoeba*. Proc Natl Acad Sci U S A 99, 1414–1419. (2002).

10. Nikolaev, S.I. et al. The twilight of Heliozoa and rise of Rhizaria, an emerging supergroup of amoeboid eukaryotes. Proc Natl Acad Sci U S A 101, 8066–8071. (2004).

11. Burki, F. et al. Phylogenomics reshuffles the eukaryotic supergroups. PLoS One. 2, e790. (2007).

12. Burki, F. et al. Large-scale phylogenomic analyses reveal that two enigmatic protist lineages, Telonemia and Centroheliozoa, are related to photosynthetic chromalveolates. Genome Biol Evol 2009, 231–238 (2009).

13. Burki, F., Okamoto, N., Pombert, J.F. & Keeling, P.J. The evolutionary history of haptophytes and cryptophytes: phylogenomic evidence for separate origins. Proc Biol Sci 279, 2246–2254 (2012).

14. Brown, M.W. et al. Phylogenomics demonstrates that breviate flagellates are related to opisthokonts and apusomonads. Proc Biol Sci 280, 20131755 (2013).

15. Brown, M.W. et al. Phylogenomics places orphan protistan lineages in a novel eukaryotic super-group. Genome Biol Evol 10, 427–433 (2018).

16. Torruella, G. et al. Global transcriptome analysis of the aphelid *Paraphelidium tribonemae* supports the phagotrophic origin of fungi. Commun Biol 1, 231 (2018).

17. Burki, F., Roger, A.J., Brown, M.W. & Simpson, A.G.B. The new tree of eukaryotes. Trends Ecol Evol 35, 43–55 (2020).

18. Simpson, A.G. & Roger, A.J. The real’kingdoms’ of eukaryotes. Curr Biol 14, R693–6 (2004).

19. Rodríguez-Ezpeleta, N. et al. Monophyly of primary photosynthetic eukaryotes: green plants, red algae, and glaucophytes. Curr Biol 15, 1325–1330. (2005).

20. Hampl, V. et al. Phylogenomic analyses support the monophyly of Excavata and resolve relationships among eukaryotic “supergroups”. Proc Natl Acad Sci U S A. 106, 3859–3864. (2009).

21. Burki, F. et al. Untangling the early diversification of eukaryotes: a phylogenomic study of the evolutionary origins of Centrohelida, Haptophyta and Cryptista. Proc Biol Sci 283, 1823 (2016).

22. Blaz, J. et al. One high quality genome and two transcriptome datasets for new species of *Mantamonas*, a deep-branching eukaryote clade. Sci Data 10, 603 (2023).

23. Tikhonenkov, D.V. et al. Microbial predators form a new supergroup of eukaryotes. Nature 612, 714–719 (2022).

24. Burki, F., Shalchian-Tabrizi, K. & Pawlowski, J. Phylogenomics reveals a new’megagroup’ including most photosynthetic eukaryotes. Biol Lett 4, 366–369 (2008).

25. Derelle, R. et al. Bacterial proteins pinpoint a single eukaryotic root. Proc Natl Acad Sci U S A 112, E693–699 (2015).

26. Simão, F.A., Waterhouse, R.M., Ioannidis, P., Kriventseva, E.V. & Zdobnov, E.M. BUSCO: assessing genome assembly and annotation completeness with single-copy orthologs. Bioinformatics 31, 3210–3212 (2015).

27. Sahbou, A.E., Iraqi, D., Mentag, R. & Khayi, S. BuscoPhylo: a webserver for Busco-based phylogenomic analysis for non-specialists. Sci Rep 12, 17352 (2022).

28. Timilsena, P.R. et al. Phylogenomic resolution of order-and family-level monocot relationships using 602 single-copy nuclear genes and 1375 BUSCO genes. Front Plant Sci 13, 876779 (2022).

29. Van Damme, K., Cornetti, L., Fields, P.D. & Ebert, D. Whole-genome phylogenetic reconstruction as a powerful tool to reveal homoplasy and ancient rapid radiation in waterflea evolution. Syst Biol 71, 777–787 (2022).

30. Strassert, J.F.H., Irisarri, I., Williams, T.A. & Burki, F. A molecular timescale for eukaryote evolution with implications for the origin of red algal-derived plastids. Nat Commun 12, 1879 (2021).

31. Petitjean, C., Deschamps, P., Lopez-Garcia, P., Moreira, D. & Brochier-Armanet, C. Extending the conserved phylogenetic core of archaea disentangles the evolution of the third domain of life. Mol Biol Evol 32, 1242–1254 (2015).

32. Eme, L. et al. Inference and reconstruction of the heimdallarchaeial ancestry of eukaryotes. Nature 618, 992–999 (2023).

33. Baker, B.A. et al. Expanded phylogeny of extremely halophilic archaea shows multiple independent adaptations to hypersaline environments. Nat Microbiol 9, 964–975 (2024).

34. Richter, D.J., et al. EukProt: A database of genome-scale predicted proteins across the diversity of eukaryotes Peer Comm J 2, e56 (2022).

35. Banos, H. et al. GTRpmix: A linked general time-reversible model for profile mixture models. Mol Biol Evol 41, msae174 (2024).

36. Eglit, Y., et al. *Meteora sporadica*, a protist with incredible cell architecture, is related to Hemimastigophora. Curr Biol 34, 451–459.e6 (2024).

37. Yazaki, E., Shiratori, T. & Inagaki, Y. Protists with uncertain phylogenetic affiliations for resolving the deep tree of eukaryotes. Microorganisms 13, 1926 (2025).

38. Zlatogursky, V. et al. Phylogenetic position and mitochondrial genome evolution of “orphan” eukaryotic lineages. iScience 28, 113184 (2025).

39. Cavalier-Smith, T. Early evolution of eukaryote feeding modes, cell structural diversity, and classification of the protozoan phyla Loukozoa, Sulcozoa, and Choanozoa. Eur J Protistol 49, 115–178 (2013).

40. Rodríguez-Ezpeleta, N. et al. Detecting and overcoming systematic errors in genome-scale phylogenies. Syst Biol 56, 389–399 (2007).

41. Lartillot, N. & Philippe, H. A Bayesian mixture model for across-site heterogeneities in the amino-acid replacement process. Mol Biol Evol 21, 1095–1109 (2004).

42. Harada, R. et al. Encyclopedia of family A DNA polymerases localized in organelles: Evolutionary contribution of bacteria including the proto-mitochondrion. Mol Biol Evol. 41, msae014 (2024).

43. Philippe, H., Delsuc, F., Brinkmann, H. & Lartillot, N. Phylogenomics. Annu Rev Ecol Syst 36, 541–562 (2005).

44. Lartillot, N., Brinkmann, H. & Philippe, H. Suppression of long-branch attraction artefacts in the animal phylogeny using a site-heterogeneous model. BMC Evol Biol 7 Suppl 1, S4 (2007).

45. Strassert, J.F.H., Jamy, M., Mylnikov, A.P., Tikhonenkov, D.V. & Burki, F. New phylogenomic aof the enigmatic phylum Telonemia further resolves the eukaryote Tree of Life. Mol Biol Evol 36, 757–765 (2019).

46. Yazaki, E. et al. The closest lineage of Archaeplastida is revealed by phylogenomics analyses that include *Microheliella maris*. Open Biol 12, 210376 (2022).

47. Bjornson, S., Lax, G., Okamoto, N. & Keeling, P.J. Phylogenomic analysis of deep-branching telonemid. Genome Biol Evol 17, evaf202 (2025).

48. Seo, T.K. Calculating bootstrap probabilities of phylogeny using multilocus sequence data. Mol Biol Evol 25, 960–971 (2008).

49. Cavalier-Smith, T., Chao, E.E. & Lewis, R. Multiple origins of Heliozoa from flagellate ancestors: New cryptist subphylum Corbihelia, superclass Corbistoma, and monophyly of Haptista, Cryptista, Hacrobia and Chromista. Mol Phylogenet Evol 93, 331–362 (2015).

50. Anderson, O.R. Cytoplasmic fine structure of nassellarian Radiolaria. Mar Micropaleontol 2, 251–264 (1977).

51. Eme, L., Sharpe, S.C., Brown, M.W. & Roger, A.J. On the age of eukaryotes: evaluating evidence from fossils and molecular clocks. Cold Spring Harb Perspect Biol 6, 8 (2014).

52. Bodył, A., Stiller, J.W. & Mackiewicz, P. Chromalveolate plastids: direct descent or multiple endosymbioses? Trends Ecol Evol. 24, 119–121 (2009).

53. Ponce-Toledo, R.I., Moreira, D., López-García, P. & Deschamps, P. Molecular phylogeny of the SELMA translocation machinery recounts the evolution of complex photosynthetic eukaryotes. Mol Biol Evol 42, msaf167 (2025).

54. Terpis, K.X. et al. Multiple plastid losses within photosynthetic stramenopiles revealed by comprehensive phylogenomics. Curr Biol 35, 483–499.e8 (2025).

55. Saville-Kent, W. *A manual of the Infusoria: including a description of all known flagellate, ciliate and tentaculiferous Protozoa, British and foreign, and an account of the organisation and affinities of the sponges*, (David Bogue, London, 1880).

56. O’Kelly, C.J. & Nerad, T.A. *Malawimonas jakobiformis* n. gen., n. sp. (Malawimonadidae n. fam.): A *Jakoba*-like heterotrophic nanoflagellate with discoidal mitochondrial cristae. J Euk Mic 46, 522–531 (1999).

57. Hehenberger, E. et al. Novel predators reshape holozoan phylogeny and reveal the presence of a two-component signaling system in the ancestor of animals. Curr Biol 27, 2043–2050.e6 (2017).

58. Leonard, G. et al. Comparative genomic analysis of the’pseudofungus’ *Hyphochytrium catenoides*. Open Biol 8, 170184 (2018).

59. Tikhonenkov, D.V. et al. New lineage of microbial predators adds complexity to reconstructing the evolutionary origin of animals. Curr Biol 30, 4500–4509.e5 (2020).

60. Cho, A. et al. Monophyly of diverse Bigyromonadea and their impact on phylogenomic relationships within stramenopiles. Mol Phylogenet Evol 171, 107468 (2022).

61. Galindo, L.J., Prokina, K., Torruella, G., López-García, P. & Moreira, D. Maturases and group II introns in the mitochondrial genomes of the deepest jakobid branch. Genome Biol Evol 15, evad058 (2023).

62. Li, W. & Godzik, A. Cd-hit: a fast program for clustering and comparing large sets of protein or nucleotide sequences. Bioinformatics 22, 1658–1659 (2006).

63. Hernández-Plaza, A. et al. eggNOG 6.0: enabling comparative genomics across 12 535 organisms. Nucleic Acids Res 51, D389–d394 (2023).

64. Buchfink, B., Reuter, K. & Drost, H.G. Sensitive protein alignments at tree-of-life scale using DIAMOND. Nat Methods 18, 366–368 (2021).

65. Eddy, S.R. Accelerated profile HMM searches. PLoS Comput Biol 7, e1002195 (2011).

66. Katoh, K. & Standley, D.M. MAFFT multiple sequence alignment software version 7: improvements in performance and usability. Mol Biol Evol 30, 772–780 (2013).

67. Criscuolo, A. & Gribaldo, S. BMGE (Block Mapping and Gathering with Entropy): a new software for selection of phylogenetic informative regions from multiple sequence alignments. BMC Evol Biol 10, 1471–2148 (2010).

68. Minh, B.Q. et al. IQ-TREE 2: New models and efficient methods for phylogenetic inference in the genomic era. Mol Biol Evol 37, 1530–1534 (2020).

69. Huerta-Cepas, J., Serra, F. & Bork, P. ETE 3: Reconstruction, analysis, and visualization of phylogenomic data. Mol Biol Evol 33, 1635–1638 (2016).

70. Camacho, C. et al. BLAST+: architecture and applications. BMC Bioinformatics 10, 421 (2009).

71. Cantalapiedra, C.P., Hernández-Plaza, A., Letunic, I., Bork, P. & Huerta-Cepas, J. eggNOG-mapper v2: Functional annotation, orthology assignments, and domain prediction at the metagenomic scale. Mol Biol Evol 38, 5825–5829 (2021).

72. Huerta-Cepas, J. et al. eggNOG 5.0: a hierarchical, functionally and phylogenetically annotated orthology resource based on 5090 organisms and 2502 viruses. Nucleic Acids Res 47, D309–D314 (2019).

73. Wang, H.-C., Minh, B.Q., Susko, E. & Roger, A.J. Modeling site heterogeneity with Posterior Mean Site Frequency Profiles accelerates accurate phylogenomic estimation. Syst Biol 67, 216–235 (2018).

74. Lartillot, N., Rodrigue, N., Stubbs, D. & Richer, J. PhyloBayes MPI: phylogenetic reconstruction with infinite mixtures of profiles in a parallel environment. Syst Biol 62, 611–615 (2013).

